# Dose, temperature and formulation shape *Metarhizium anisopliae* virulence against the oriental fruit fly: lessons for improving on-target control strategies

**DOI:** 10.1101/2023.12.14.571642

**Authors:** Anaïs Chailleux, Oumou Noumou Coulibaly, Babacar Diouf, Samba Diop, Ahmad Sohel, Thierry Brévault

**Affiliations:** CIRAD, UPR HortSys, Biopass, Centre de recherche ISRA-IRD, Dakar, Senegal; CIRAD, Univ Montpellier, Montpellier, France; UCAD, ED-SEV, Dakar, Senegal; Joint FAO, IAEA Division of Nuclear Techniques in Food and Agriculture IPCL, Seibersdorf, Austria; CIRAD, UPR AIDA, Biopass, Centre de recherche ISRA-IRD, Dakar, Senegal

**Keywords:** *Metarhizium anisopliae*, *Bactrocera dorsalis*, entomovector technology, auto-dissemination, boosted SIT, pest management

## Abstract

Entomopathogenic fungi are a promising tool for the biological control of crop pests provided low or no impact on non-target organisms. Selection for host specificity as well as on-target applications open new avenues for more sustainable strategies for pest management. Isolates of *Metarhizium anisopliae* (Metschn.) Sorokin have been identified as promising for developing innovative entomovectoring-based strategies for the control of the oriental fruit fly, *Bactrocera dorsalis* (Hendel) (Diptera: Tephritidae), in Africa. To be effective, this technology requires high strain virulence at a low number of spores, but sufficient incubation time to allow transmission to wild conspecifics. This depends on trophic interactions between the host and the pathogen, which are mediated by abiotic factors.

In the present study, we investigated the virulence of the Met69 strain against adult flies, depending on the inoculation dose, air temperature and formulation. High pathogenicity was observed at very low inoculation doses (LT50 of 4.85 days with 6100 spores per fly) independently of fly sex. Virulence increased with spore load in a tight range (5600 and 6100 spores per fly) and with air temperature observed in the field (20-28°C). Unexpectedly, corn starch used as an adjuvant to increase the carrying capacity of insects decreased the virulence of the pathogen.

The results will help improve area-wide control strategies based on the contamination of wild flies through auto-inoculation devices or interactions with released mass-reared sterile males coated with fungal spores. Furthermore, the study proposes an approach for calibrating area-wide control strategies, taking into account both the insect and pathogen bioecology and the environment in which they evolve.

**Author roles:** **Anaïs Chailleux:** Conceptualization, Funding acquisition, Methodology, Data curation, Writing – original draft. **Oumou Noumou Coulibaly:** Investigation, Writing – original draft. **Babacar Diouf:** Investigation, Visualization. **Samba Diop:** Investigation. **Ahmad Sohel:** Resources, Writing – review & editing. **Thierry Brévault:** Conceptualization, Funding acquisition, Project administration, Writing – review & editing.

## Introduction

Entomopathogens are among the main natural enemies of arthropod pests in tropical agroecosystems (Meyling and Eilenberg 2007, Hawkins et al. 1997). They include bacteria, fungi, protozoa, nematodes or viruses that can act as biocontrol agents. Their increasing use for crop protection is encouraged by a general trend towards the ‘zero-pesticide’ farming challenge and the agroecological transition of cropping systems. The use of entomopathogenic microorganisms presents several advantages such as safety for humans, medium to high specificity, and low risk of resistance evolution (Lacey et al. 2001, Singh et al. 2017). In addition, strain selection, as well as innovative formulations and application methods, can act as levers to increase the specificity and virulence to the target pest (Lacey 2001).

One of the main expected improvements of the use of entomopathogenic microorganisms for pest management is on-target application to reduce unintentional impact on non-target organisms and thus arthropod biodiversity (Leite et al 2022). In Africa, a strain of *Metarhizium anisopliae* (Metschn.) Sorokin (Hypocreales: Clavicipitaceae) has been identified as promising for the control of the oriental fruit fly, *Bactrocera dorsalis* (Hendel) (Diptera: Tephritidae), particularly through soil application in mango orchards (Ekesi et al. 2011). One issue of such treatments is that *M. anisopliae* spores, because of generally low specificity, threaten non-target arthropod species including beneficials (Thungrabeab and Tongma 2007, Zimmermann 2007). Therefore, the design of innovative based, for example, entomovectoring such as auto-inoculation of wild flies in the field (Faye et al. 2021, Stafford 2017) or the release of mass-reared sterile males (in the framework of Sterile Insect Technique [SIT] programs,) as vectors of micro-doses of biocides to wild flies of the same species (Bouyer and Lefrançois 2014, Diouf et al. 2022), open new avenues for more sustainable strategies for fruit fly management. The latter technology, named ‘boosted SIT’, has shown some potential in coffee-growing areas in Guatemala where the release of *C. capitata* sterile males coated with fungal spores of *Beauveria bassiana* resulted to spore transmission to 44% of the captured wild males (Flores et al. 2013).

To be effective, this technology requires high strain virulence to kill the contaminated wild individuals with a low number of spores, but with a sufficient incubation period to allow sufficient time for transmission from mass-reared insects to wild conspecifics. This depends on the trophic interactions between the host and the pathogen, which are mediated by abiotic factors of the environment where the strategy will take place. Until now, no information is available on the virulence at micro-doses of *M. anisopliae* spores on *B. dorsalis* adults. Most studies on the virulence of entomopathogenic fungi have considered doses for soil application to control larvae and pupae (Abdellah et al. 2020, Tora & Azerefegn et al. 2021). Furthermore, the impact of abiotic conditions on virulence of this entomopathogenic fungus is poorly known. Temperature optimum is variable among fungus species and strains (Thungrabeab et al 2006, Filotas et al 2006, Quesada-Moraga 2006a) and, unexpectedly, is not necessarily linked to their geographical origin (Meyling & Eilenberg 2007, Devi et al. 2005, López Plantey et al. 2019). Virulence is the result of a fight between the pathogen and the insect that depends on the optimum of temperature that will favor spore germination and mycelium growth (Yeo et al. 2003, Nussenbaum et al. 2013), and on the optimum temperature of the insect immune response which also depends on the immune mechanism involved (Murdock et al. 2012). Thus, mycelium development and immune response patterns observed under one set of conditions on a given host provide little basis for predicting virulence in other conditions, which is rather shaped by the fungus–insect interactions mediated by local context (James et al. 1998, Kryukov et al. 2018, Yeo et al. 2003). Lastly, pathogen virulence could be modified by adjuvants, also called ‘carrier’ or ‘diluents’ (Mommaerts et al. 2012, Rogers et al. 2014), that are added to spores to increase the carrying capacity of entomovectors (hereafter, ‘spore load’). Among them, corn starch particles (Escande 2002, Al-mazra’awi et al. 2006, Kevan et al. 2008, Smagghe et al. 2013) have the potential to increase spore load of vectors as they aggregate spores.

Another relevant mediator parameter in insect immunity is the sex of individuals, but only a few recent studies have investigated this aspect. Duneau et al. (2024) showed significant variations in the mortality induced by different strains of *M. anisopliae* in males *B. dorsalis*, but not in females that exhibited low mortality. Strains varied in their sub-lethal effects on female fecundity. This might be explained by differential responses to fungal infection between sexes, as was found for the expression profile of antimicrobial peptide genes in *Ceratitis capitata* when infested by *Purpureocillium lilacinum* (Djobbi et al. 2023). Moderator effects can also vary according to sex. Rantala et al. (2020) showed that the administration of juvenile hormone (a key regulatory molecule in the development and life cycle of insects) prolonged survival time after infection with *Metarhizium robertsii* in males but reduced survival time in females.

The objective of the present study was to evaluate the virulence of *M. anisopliae* spores (strain Met69) on *B. dorsalis* adult flies according to the inoculation dose and to the actual spore load. We also investigated the effect of formulation (adjuvant) and Senegalese seasonal temperature on the pathogen virulence. Results are discussed in the light of improvement of the entomovector technology for the sustainable management of pest.

## Material and methods

### Fungal spores

*Metarhizium anisopliae* Met69 (Real IPM Ltd, Kenya, [REAL IPM UK, 2015]) was supplied as pure spore powder. The spore powder contained 1,89 e^+10^ (±0.15 e^+10^) spores. g^-1^, spore length was 6,57 (±0.30) µm, and width was 2,46 (±0.26) µm. Its germination rate was 71.8% (± 4.0) after 24 h at 27° C, and 90.2% (±1.2) after 48 h.

### Fruits flies

The entomopathogen was tested on two distinct lab-reared populations of the oriental fruit fly, *B. dorsalis*: sterile individuals provided by the Insect Pest Control Laboratory (IPCL) of the Joint FAO/IAEA Division of Nuclear Techniques in Food and Agriculture (Seibersdorf, Austria), and local fertile individuals produced in our Laboratory (Biopass, Dakar, Senegal). Sterile flies were shipped as pupae by air cargo companies (3–4-day journey) as described by Chailleux *et al*. (2021). Local fertile flies were obtained by collecting and incubating mangoes from the Niayes area in Senegal in 2019 with yearly addition of wild flies (∼100 individuals/year). They were reared in the laboratory in 45 × 45 × 45 cm insect cages (BugDorm, Taiwan) at 27°C and 70% RH. Water, sugar, and yeast extract powder (Alfa Aesar, Kandel, Germany) provided *ad libitum*. Fresh mature bananas were provided as an oviposition substrate. After 48 hours, bananas were incubated in plastic bins containing sterilized sand. Pupae were then placed in a rearing cage until adult emergence. Only healthy and sexually mature flies (10 to 14 days old) were used in the experiments.

### Dose-mortality relationship

Flies of the local fertile population were inoculated with spores using a cylindrical plastic tube (8 × 6 cm) lined with velvet containing Met69 spores. Ten flies were introduced all together into the tube and exposed to the conidia for 3 min. They were then transferred to cages for three hours to allow them time to groom without contaminating the experimental cages. A series of 10 doses ranging from 0 (control) to 6.4 × 10^8^ spores per square centimeter was tested. For each dose, 60 flies (30 fertile males and 30 fertile females) were inoculated. The use of spores per square centimeter allows the standardization of the incubation doses across studies. Indeed, the size of the tube is not of importance, as we showed that it did not impact fly inoculation if the number of spores per unit surface is respected (Appendix 1). As this design provided high mortality even for the smallest dose, we changed our inoculation method to be able to reduce the spore load on flies. To this end, a paintbrush with a reduced number of hairs (8, 4, 2, and 1 hair) was used to manually apply the spore powder to the body of the flies. The fewer hairs on the paintbrush, the fewer spores brought to the flies.

Among the 30 flies of each sex, 10 were used to count the number of spores collected by the individuals after grooming. For this purpose, flies were individually put in a tube with 1 ml of distilled water and a drop of Tween 80, then vortexed for 3 min (Appendix 2). A sample of the solution was taken for counting spores in Malassez cells under a microscope at 40x magnification. The remaining 20 flies were kept for 15 days for daily mortality monitoring (27

± 2° C). Flies were placed individually in transparent entomological boxes (3 × 8cm) with an aeration grid (Entomo-Silex, France). A mixture of yeast hydrolysate and sugar was placed inside each box to feed the flies, and cotton soaked with water was placed on the grid. Dead flies were incubated in a climate cabinet (27 ± 2° C) on a wet sponge in a Petri dish to diagnose the cause of death (check for fungal development).

### Effect of temperature on spore germination and growth

A range of temperatures (monthly average) close to field conditions was selected to be representative of the Niayes area, the main production basin of mango in North Senegal. Using a climatic chamber (I-30 VL, Percival Scientific, Inc., USA), three temperatures (mean night temperature/ mean day temperature) were tested, corresponding to months of February (17.2 / 22.8°C), May (20.7 / 24.3°C), and October (26.0 / 30.2°C) in the Niayes area, based on 2017 to 2020 data of a weather station located in Sangalkam (GPS coordinates: 14.789468, - 17.226484). February is the coldest month with low population of *B. dorsalis* (middle of the dry season), May is the month when *B. dorsalis* population starts increasing (end of the dry season), and October the hottest when *B. dorsalis* population starts decreasing (end of the rainy season). To assess the impact of day/night alternation, the average temperatures of these three months were also tested in a constant regime (20.1; 22.5; 27.7). The relative humidity was 70-80% and the photoperiod was 12/12 (D/L).

Spore germination was assessed by inoculating an SDA media (Sabouraud Dextrose Agar) in a Petri dish with a conidial suspension (concentration of 1 × 10^−5^ g.ml) made with 0.01 g of dry spore powder, one drop of Tween 80, and 10 ml of distilled water and then diluted at 1%. Petri dishes were then sealed with parafilm and incubated at the six temperatures. Spore germination was observed at 24 and 48 h, as long as the development of the mycelium allowed us to measure the evolution. The germination rate was determined by examining 100 randomly selected spores per dish using a microscope at 40x magnification. Five replicates were performed per temperature. Conidia were considered germinated when they were longer than normal conidia (Petlamul and Prasertsan, 2012). The same procedure was adopted to assess mycelial growth, but only three drops of the spore suspension were placed in the middle of the Petri dish on the SDA medium. Five replicates were performed per temperature. Fungal growth (mm) in each dish was determined by measuring the average diameter of two perpendicular lines previously drawn on the bottom of the Petri dish daily for 7 days (Membang et al., 2021).

### Effect of temperature on pathogen virulence

Both sexes of the local fertile population were tested, whereas only males (sterile males as entomovectors) of the IPCL population were tested. The same inoculation procedure of flies as described above was adopted but only with the dose of 4.0 × 10^7^spores.cm^-2^. The selected dose was informed by prior results, with the objective of achieving mortality while still providing time for transmission to conspecifics. Flies were then incubated at the same alternating temperatures and constant temperatures. Monitoring of mortality was done daily for 3 weeks. Dead flies were cleaned with alcohol 70% and incubated at the test temperatures in Petri dishes containing moistened sponge for 7 days to diagnose whether mortality was due to fungal infection.

### Effect of the adjuvant on pathogen virulence

The effect on virulence of corn starch (Maïzena) as an adjuvant to pure spore powder (1:1) was tested at the same alternating and constant temperature regimes, but only on the local fertile population. The quantity of spores in the inoculation tube was kept constant.

### Statistical analyses

All statistical analyses were performed using R software (R Core Team 2020) version 4.0.5. All data are available in a dataverse (Chailleux, 2023). Survival data were analyzed using Cox models which is adapted to truncated data (survival package [Therneau 2022]). For dose-mortality relationship the Cox model was built with either fly’ load or dose, and fly sex as explanatory variables. Correlation between tube doses and fly load was examined using the Pearson correlation (ggpubr package [Kassambara, 2020]). Lethal doses and lethal time were calculated using the probit method (ecotox package [Hlina et al. In press]). Survival graph were made using the package survminer (Kassambara et al. 2021). Temperature effect on mycelium germination and growth was analyzed using Generalized Linear Models (GLM) with month temperature, temperature regime (day and night vs constant), and elapsed time since inoculation as explanatory variables. A Poisson distribution was implemented for the germination and a gaussian one for the growth. Temperature effect was analyzed using Cox models built with month, temperature regime, sex, and sterile vs local population, as explanatory variables. Unless the quality control led by the IPCL on their flies (FAO/IAEA/USDA. 2019), what we called thereafter “population” discriminate the population effect, with the sterile population encompassing inseparably characteristics owing from fly population and sterilization process. Effect of the adjuvant was analyzed similarly but with the adjuvant presence, month temperature, temperature regime and sex as explanatory variables.

## Results

### Dose-mortality relationship

All the tested doses using the inoculation device induced high mortality among flies (between 80-100%), independently of the sex (χ^2^ = 0.806; df = 1; P = 0.369), but at variable speed depending on the dose (χ^2^ = 252.27; df = 1; P < 0.001) (figure 1). The lethal dose 50 (LD 50, lower dose to kill 50% of flies) after 7 days was 1.58 e^+5^ spores/cm^2^ of velvet.

**Figure 1.**
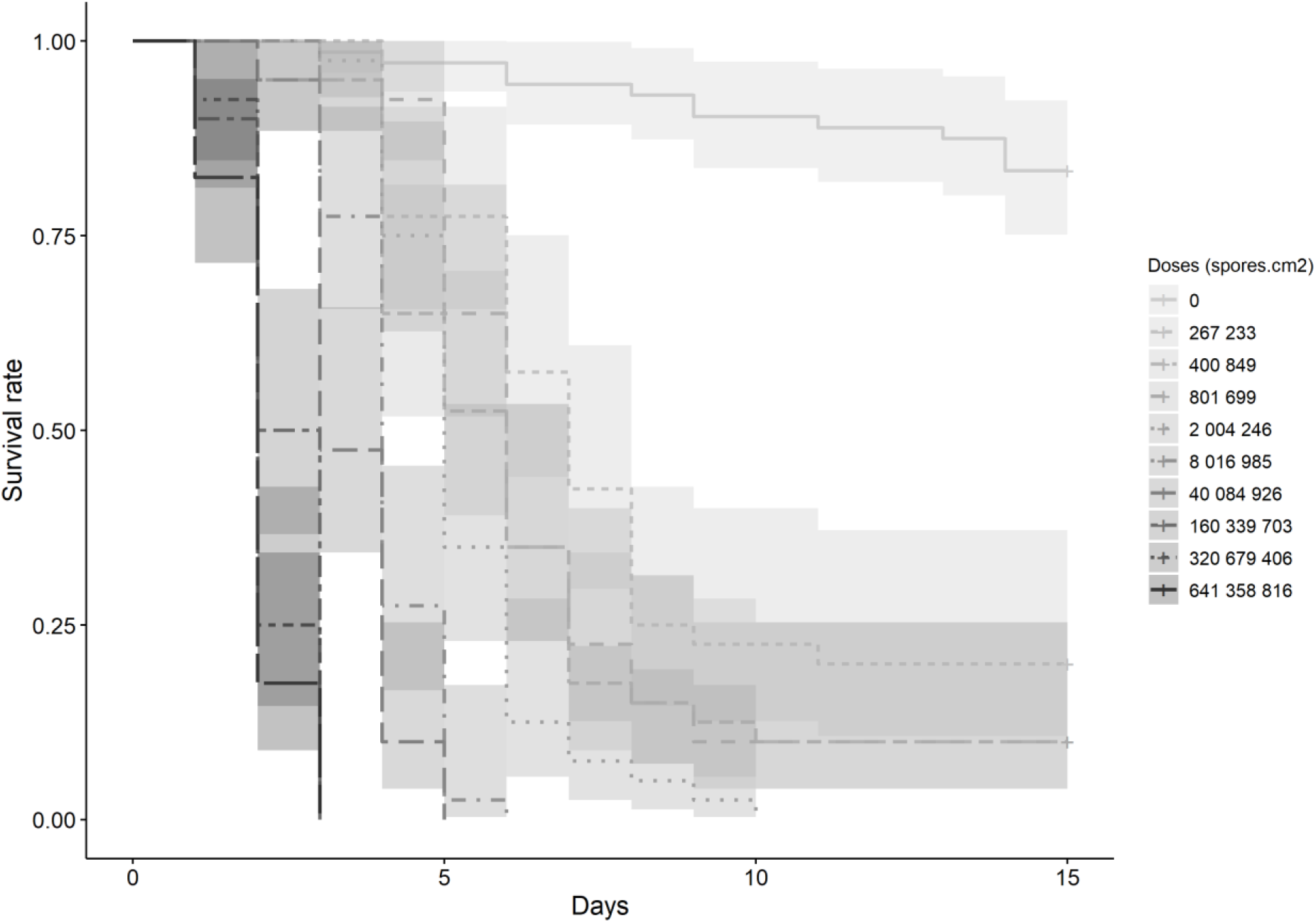
Survival rate of *B. dorsalis* exposed to a range of *M. anisopliae* (strain Met69) spore doses in the inoculation tube.

The dose threshold to obtain fast and high mortality was at the transition zone (where the slope of the curve goes from close to 1 to close to 0), between 40 084 926 and 8 016 985 spores.cm^-2^, where the LT 50 (lower number of days to kill 50% of flies) went from 3.07 to 3.63 days (figure 2). The LT 50 kept small, 5.64 days, with 2 004 246 spores.cm^-2^ but jumped to 22.9 days with 801 699 spores.cm^-2^.

**Figure 2.**
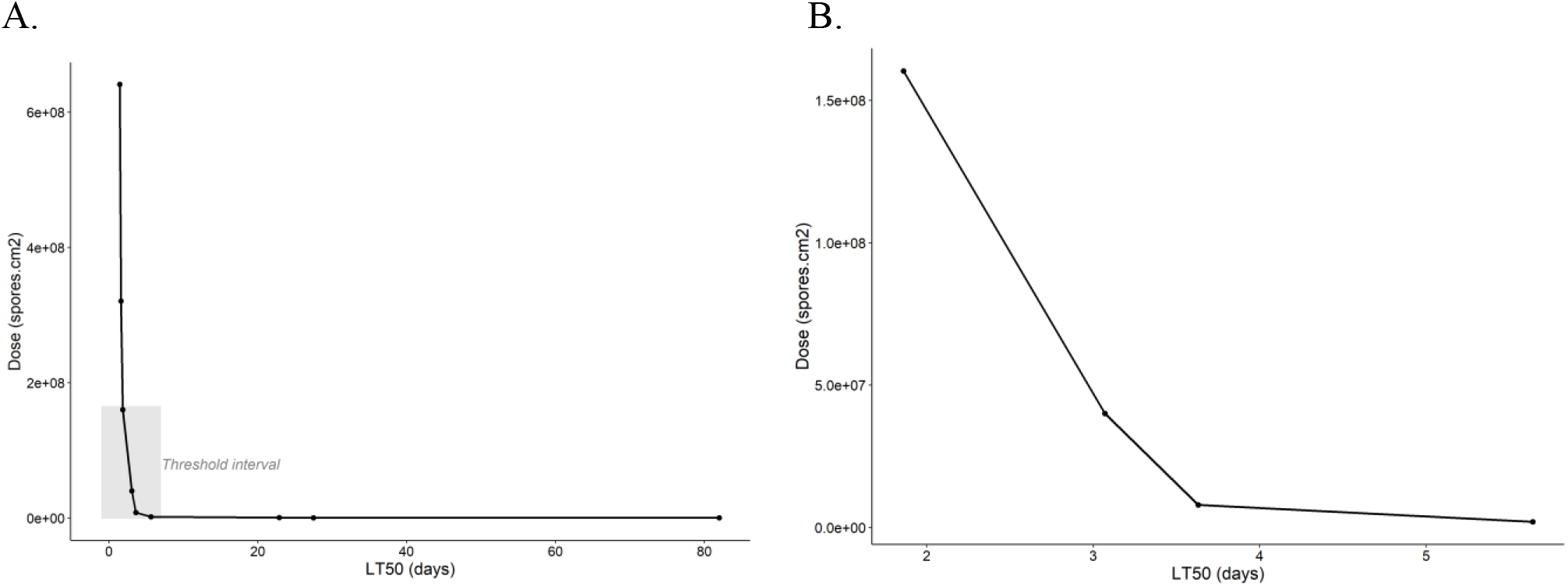
Virulence (LT50) of *M. anisopliae* strain Met69 against *Bactrocera dorsalis* flies according to inoculation dose. A. All the data set, B. Zoom on the inoculation dose threshold interval.

**Figure 3.**
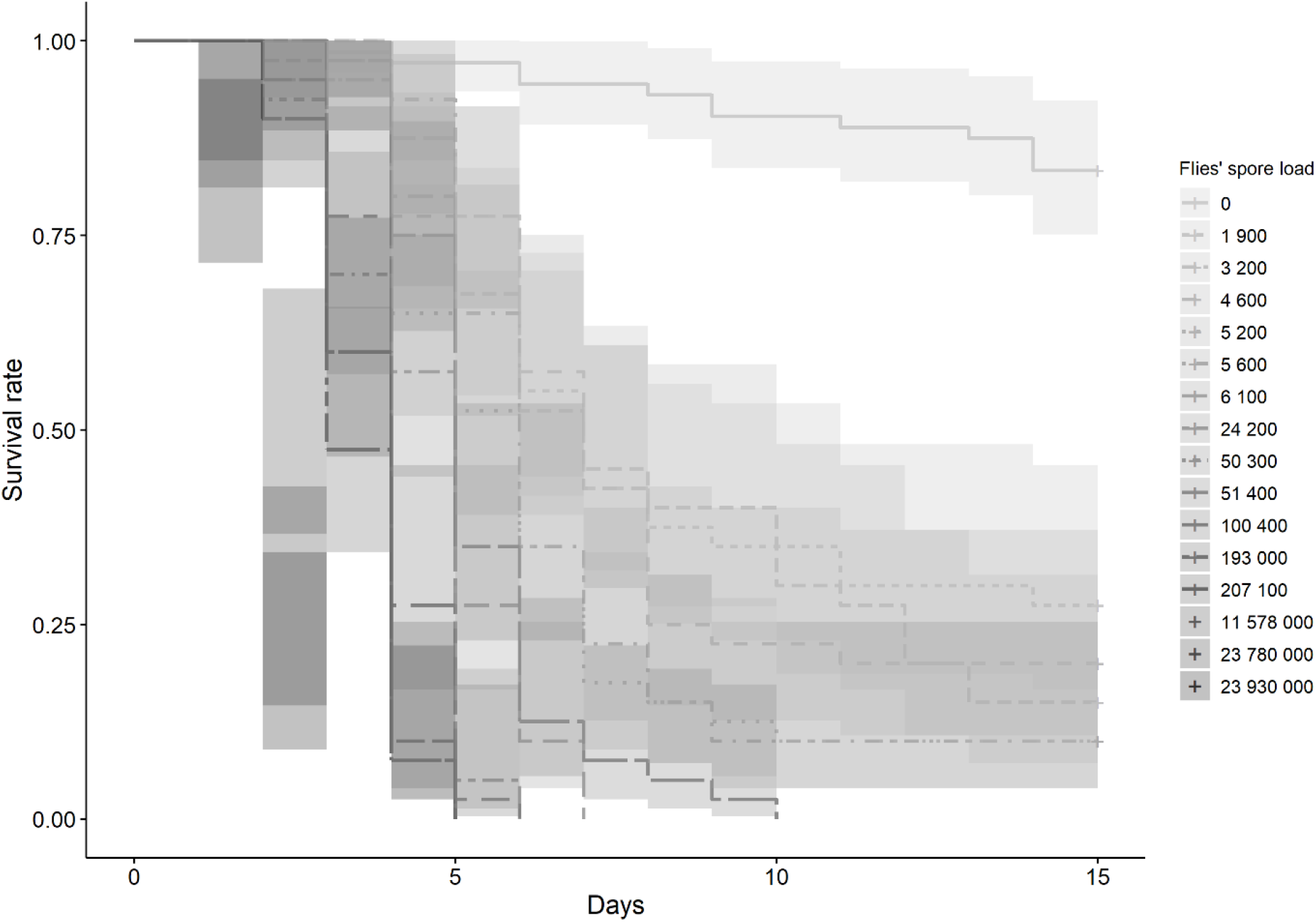
Survival rate of *B. dorsalis* exposed to a range of *M. anisopliae* (strain Met69) spore loads.

As expected, different inoculation doses using the inoculation device translated into different spore loads on flies (Pearson’s correlation: r = 0.83; P < 0.001) (Appendix 3). Data of mean spore loads obtained from the inoculation device and inoculation with paintbrush were then pooled together. Spore load significantly affected fly survival (χ^2^ = 377.682; df = 1; P < 0.001) while sex did not (χ^2^ = 0.311, df = 1; P = 0.577). LD 50 at 7 days was of 1.69 e^+3^ spores per fly.

Regarding the LT 50, spore load threshold for fast and high mortality was at the slope change, between 207 100 (paintbrush inoculation) and 11 578 000 (tube inoculation) spores per fly, where the LT 50 went from 1.86 to 3.13 days (figure 4). The LT 50 kept small, 4.85 days, with 6100 spores per fly, but jumped to 22.9 days with 5600 spores per fly.

**Figure 4.**
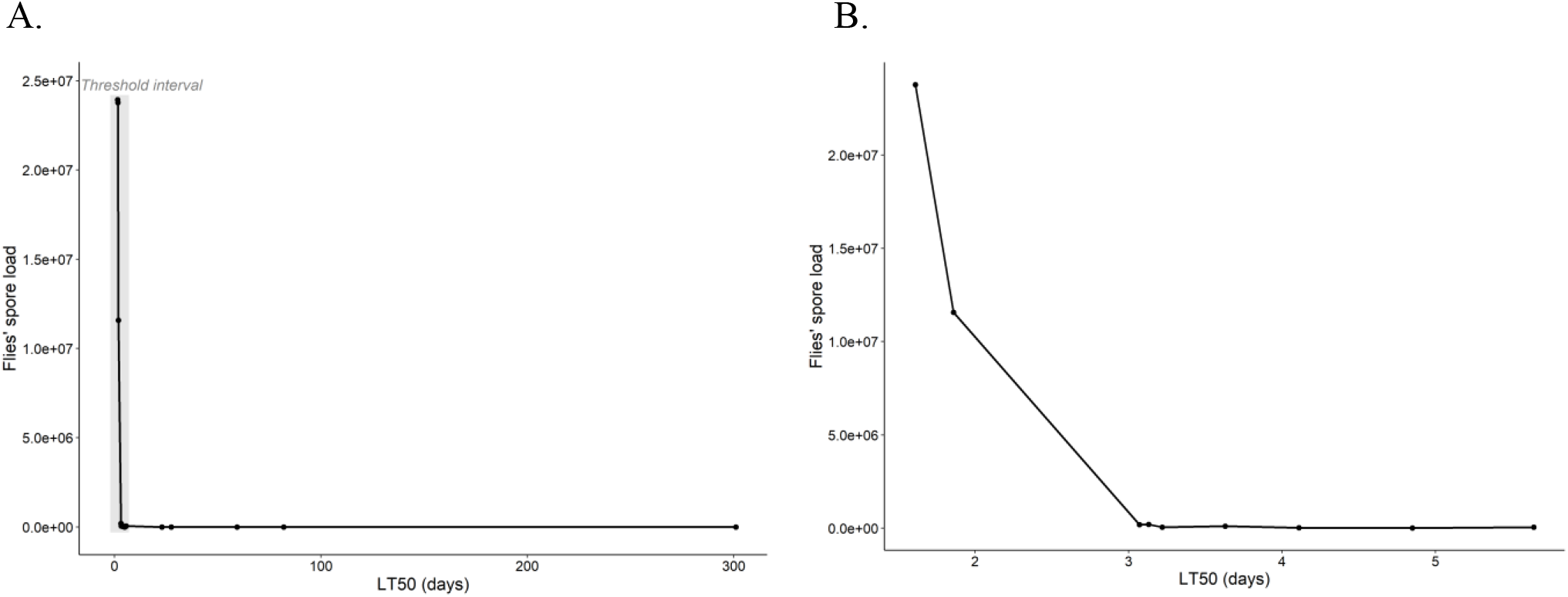
Virulence (LT50) of *M. anisopliae* strain Met69 against *Bactrocera dorsalis* flies according to spore load. A. All the data set, B. Zoom on the spore load threshold interval.

### Effect of temperature on spore germination and growth

Germination of Met69 spores was positively affected by elapsed time after inoculation (D = 8.34, df =1, 55, P =0.004) while month temperature and constant *vs* day and night regime had no effect (respectively D = 0.360, df = 2.57; P = 0.835 and D = 0.352, df = 1, 56; P = 0.553). The overall mean germination rate (±SE) after 24 h was 92.2 ± 0.9 % and 99.5 ± 0.2 % after 48 h.

Mycelium growth was affected by elapsed time after inoculation (D = 13650.8, df = 1, 205; P > 0.001) but also by month temperature (D = 4632.1, df = 2, 207; P <0.001) and temperature regime (D= 409.5, df = 1, 206; P = 0.006). The night and day temperature alternation allowed faster growth (figure 5) than constant temperature. The lowest growth was observed at the intermediate May temperature (20.7-24.3°C).

**Figure 5.**
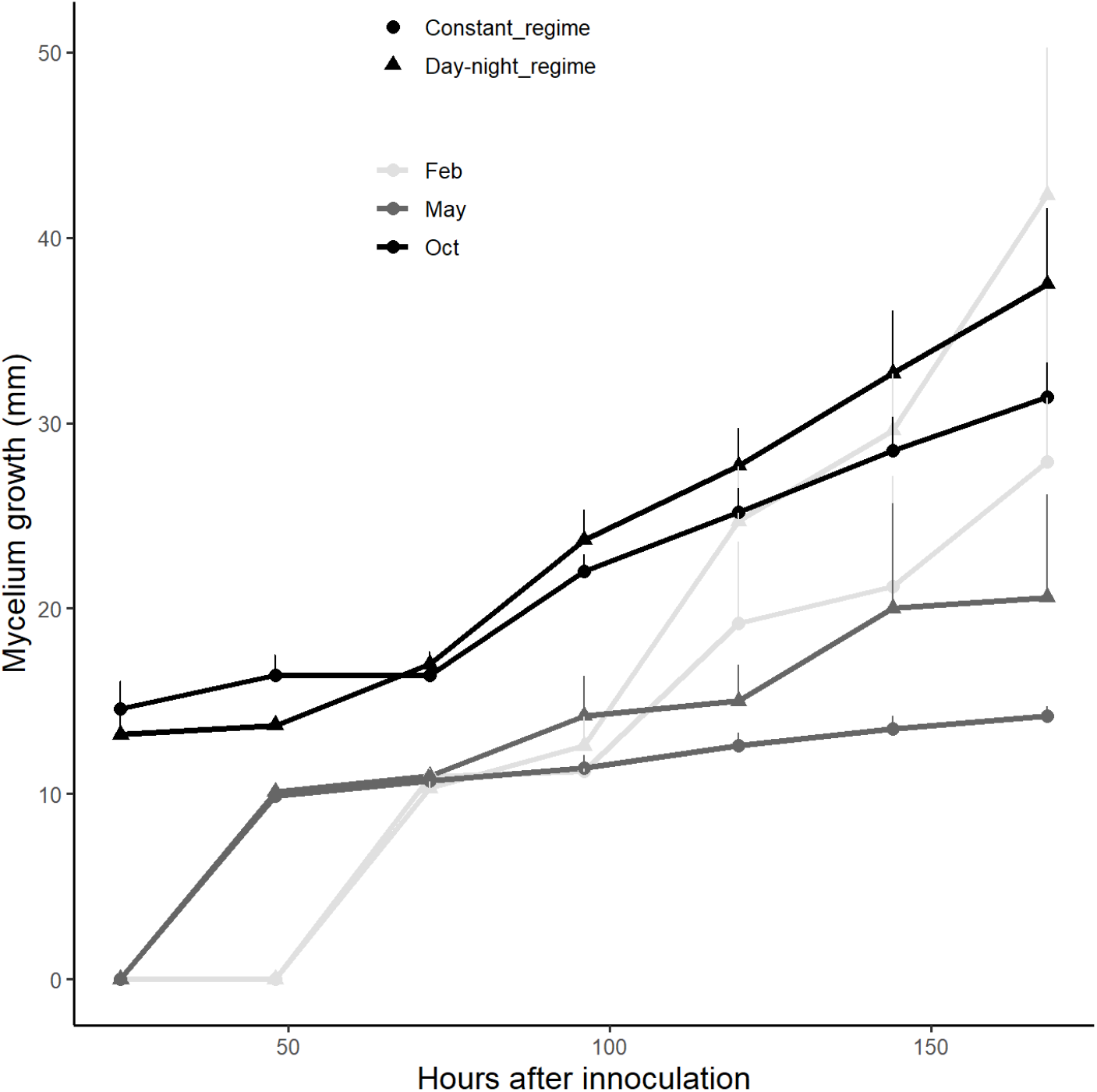
Mycelium growth according to elapsed time after fly inoculation with *Metarhiziium anisopliae* (strain Met69) at different temperatures.

### Effect of air temperature on pathogen virulence

Survival of flies was negatively affected by month temperature increase (table 1, figures 6 and 7), but the interaction between month temperature and population reveals that month temperatures effect differed between sterile and fertile males. Survival was not affected by constant *vs* day and night temperature regime (table 1). Survival of flies was negatively affected by the spore inoculation (whether or not the flies were inoculated), but the significant interaction between month and inoculation revealed that fungus virulence depended on month temperatures. Sex was the sole factor with a significant main effect without any interaction, indicating that its effect was independent of the other factors.

**Table 1.** Effect of spore inoculation with *Metarhizium anisopliae* (strain Met69), month temperature and regime, sex and population on survival rate of *Bactrocera dorsalis* flies. Cox model analysis. Significances are considered at P < 0.05.

| Variable | $\chi^2$ | Df | P value |
| --- | --- | --- | --- |
| Spore inoculation (yes/no) | 958.95 | 1 | **<0.001** |
| Month Temperature | 21.12 | 2 | **<0.001** |
| Temperature regime | 0.43 | 1 | 0.510 |
| Sex (M/F) | 5.84 | 1 | **0.015** |
| Population (sterile vs fertile males) | 38.84 | 1 | **<0.001** |
| Month: ND-C | 1.95 | 2 | 0.377 |
| Month temperature:Spore inoculation | 97.23 | 2 | **<0.001** |
| Temperature regime:Spore inoculation | 0.32 | 1 | 0.570 |
| Month temperature:Sex | 4.81 | 2 | 0.090 |
| Temperature regime:Sex | 1.43 | 1 | 0.232 |
| Spore inoculation:Sex | 0.46 | 1 | 0.497 |
| Month temperature:Fly population | 7.14 | 2 | **0.028** |
| Temperature regime: Fly population | 1.66 | 1 | 0.197 |
| Spore inoculation: Fly population | 0.06 | 1 | 0.804 |
| Sex: Fly population | 0.00 | 0 | 1.00 |

**Figure 6.**
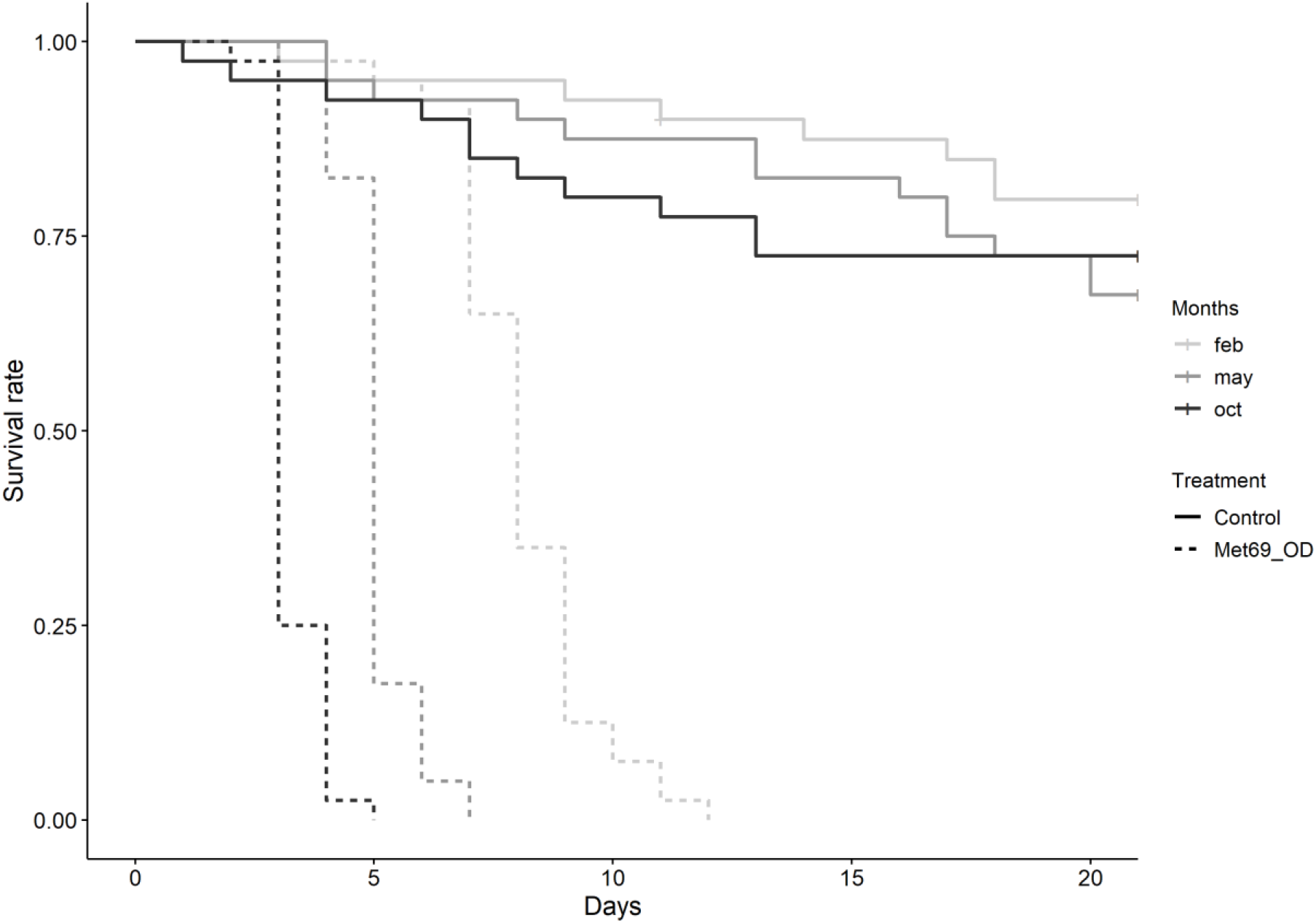
Survival rate of *Bactrocera dorsalis* females according to month temperature and inoculation with *Metarhizium anisopliae* spores (strain Met69).

**Figure 7.**
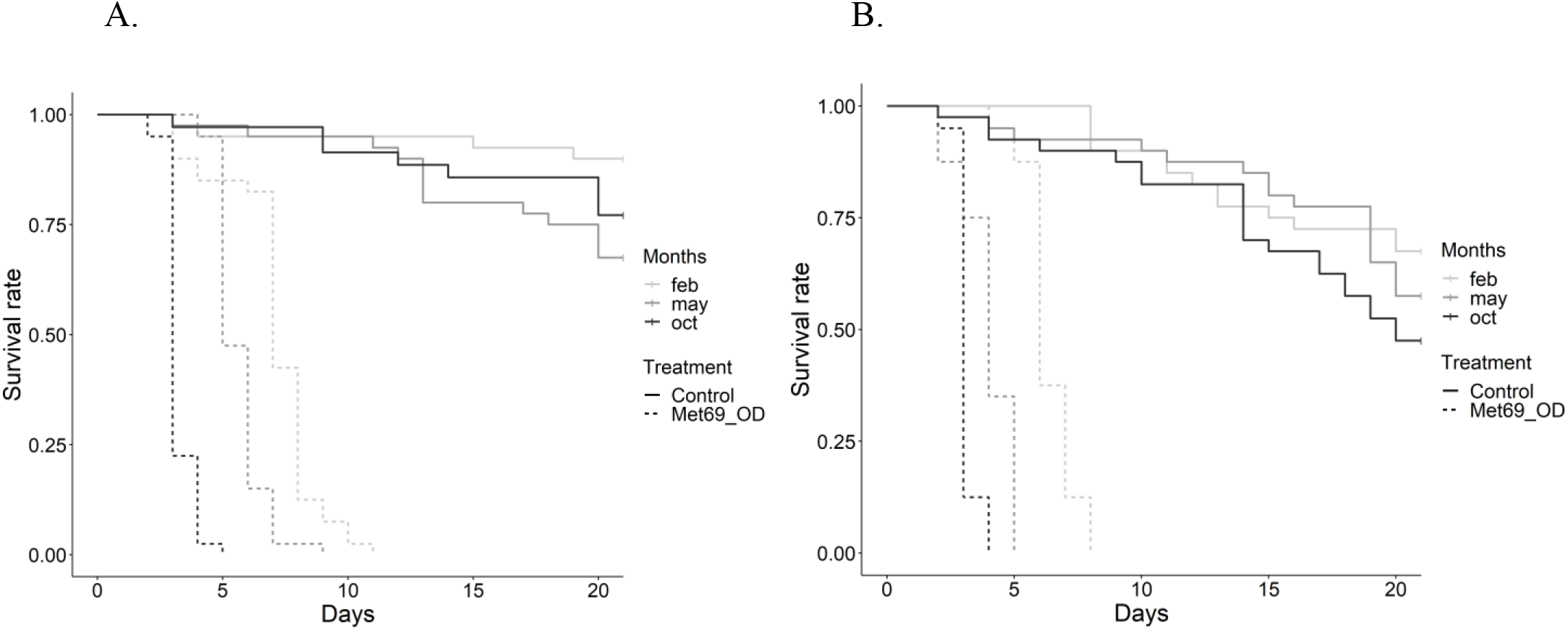
Survival rate of *Bactrocera dorsalis* males according to month temperature and inoculation with *Metarhizium anisopliae* spores (strain Met69). Fly population: (A) fertile, (B) sterile.

Among the fertile population, females tended to survive less than males. Across populations, sterile males survived less than fertile ones and this effect became more pronounced when they were inoculated (table 2). In fact, when inoculated, no fly survived up to 20 days (table 2). However, the 5- and 10-days survival rates illustrated the effect of the tested variables. After 5 days, the best survival was observed in females at the coldest temperature (February), and the worst was observed in sterile males at the highest temperature. After 10 days, only individuals of the fertile population survived at the coldest month temperature (October).

**Table 2.** Survival rate (%) of *Bactrcera dorsalis* fruit flies after 5, 10, and 20 days according to inoculation with *Metarhizium anisopliae* spores (strain Met69), temperature (month), sex and fly population.

| Inoculation | Temperature | Sex | Fly population | ≤ 5 j | ≤ 10 j | ≤ 20 j |
| --- | --- | --- | --- | --- | --- | --- |
| Control | February | F | fertile | 95 | 92.5 | 77.5 |
|  |  | M | fertile | 95 | 95 | 90 |
|  |  | M | sterile | 100 | 90 | 72.5 |
|  | May | F | fertile | 95 | 87.5 | 72.5 |
|  |  | M | fertile | 97.5 | 95 | 75 |
|  |  | M | sterile | 95 | 92.5 | 65 |
|  | October | F | fertile | 92.5 | 80 | 72.5 |
|  |  | M | fertile | 97.5 | 92.5 | 87.5 |
|  |  | M | sterile | 92.5 | 87.5 | 52.5 |
| Met69 | February | F | fertile | 97.5 | 12.5 | 0 |
|  |  | M | fertile | 85 | 7.5 | 0 |
|  |  | M | sterile | 95 | 0 | 0 |
|  | May | F | fertile | 82.5 | 0 | 0 |
|  |  | M | fertile | 95 | 0 | 0 |
|  |  | M | sterile | 35 | 0 | 0 |
|  | October | F | fertile | 2.5 | 0 | 0 |
|  |  | M | fertile | 2.5 | 0 | 0 |
|  |  | M | sterile | 0 | 0 | 0 |

### Effect of the adjuvant on pathogen virulence

The survival rate of inoculated flies significantly increased in the presence of the corn starch adjuvant, but significant interaction between adjuvant and month temperature (table 3) revealed that adjuvant effect decreased with temperature (figure 8). Significant interaction between adjuvant and temperature regime (table 3) indicated that adjuvant effect was greater under day-night regime than under constant regime (figure 8). Here, no effect of sex on the survival rate was recorded.

**Table 3.** Effect of adjuvant, temperature, and sex on the survival rate of *Bactrocera dorsalis* flies inoculated with *Metarhizium anisopliae* (strain Met69) spores. Cox model analysis. Significances are considered at P < 0.05.

| Variable | $\chi^2$ | Df | P value |
| --- | --- | --- | --- |
| Adjuvant (yes/no) | 27.47 | 1 | <0.001 |
| Month temperature | 356.05 | 2 | <0.001 |
| Temperature regime | 40.12 | 1 | <0.001 |
| Sex (M/F) | 0.08 | 1 | 0.776 |
| Adjuvant:Month Temperature | 25.54 | 2 | <0.001 |
| Adjuvant:Temperature regime | 34.57 | 1 | <0.001 |
| Month temperature:Temperature regime | 15.46 | 2 | <0.001 |
| Adjuvant:Sex | 0.05 | 1 | 0.828 |
| Month temperature:Sex | 2.15 | 2 | 0.341 |
| Temperature regime:Sex | 0.03 | 1 | 0.860 |

**Figure 8.**
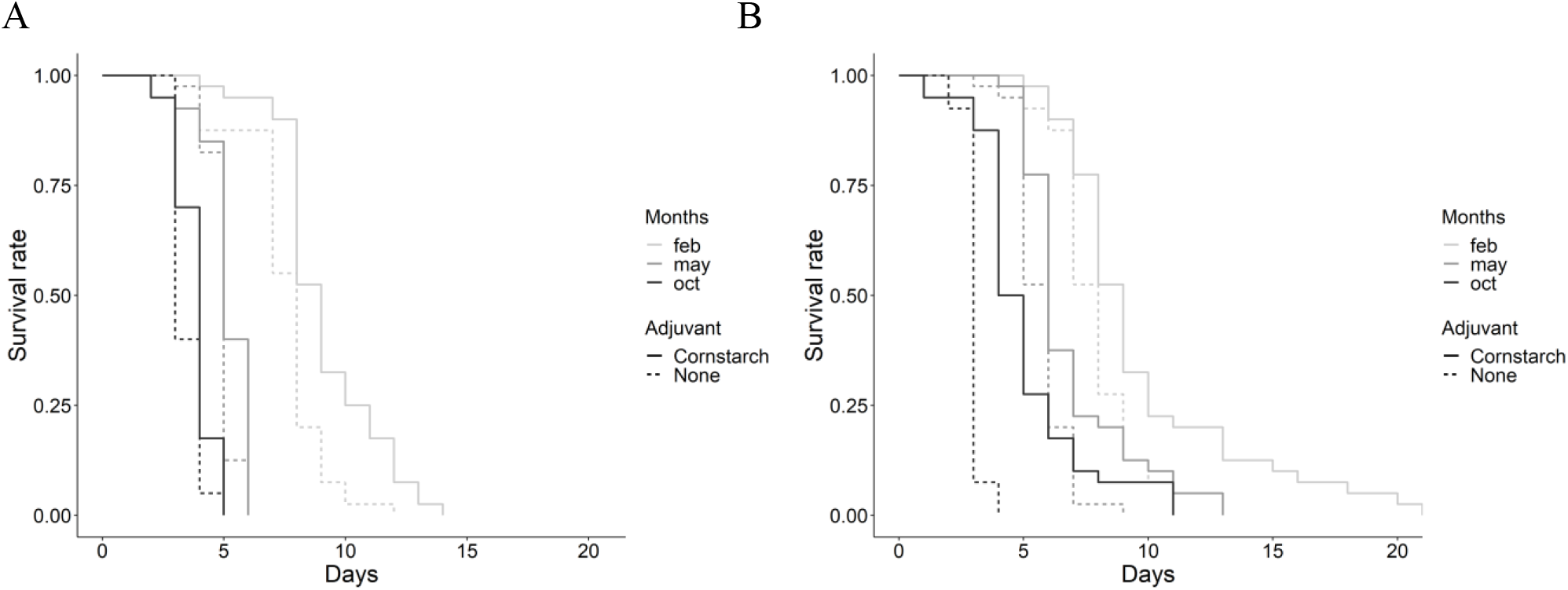
Survival rate of *Bactrocera dorsalis* flies inoculated with *Metarhizium anisopliae* (strain Met69) spores according to the presence of an adjuvant (corn starch) and temperature (A, constant regime and B, day-night variation).

## Discussion

Bioassays with *Metarhizium anisopliae* (strain Met69) against the oriental fruit fly showed that virulence increased with inoculation dose and spore load, even if high pathogenicity was observed at very low doses for both fly sexes. The pathogen virulence also increased with air temperature within the range of field temperatures tested. The slowest induction of mortality was observed for the coldest month temperature in the Niayes area in Senegal, i.e., February. Unexpectedly, corn starch used as an adjuvant increased the delay to death, which is interesting for conspecific transmission in the perspective of improving the entomovector technology.

Generally, studies of the dose-mortality relationship in insects rely on the application of spore solutions or spore feeding by adults, with a lack of information of the actual number of spores ingested or loaded by individuals (Quesada-Moraga et al. 2006b, Beris et al. 2013, Wang et al. 2021), which makes comparisons with our results difficult. Moreover, studies investigating the pathogenicity of fungal spores on fruit fly adults have not tackle the dose effect (Dimbi et al. 2003, Onsongo et al. 2022). Here, we investigated the dose-mortality relationship to identify two thresholds for both inoculation dose and spore load: a ‘minimum’ to induce mortality, and a ‘maximum’ (saturation rate) above which no more effect on the proportion of dead individuals or on the speed of the death is observed. Contrary to our expectations, the area between those two thresholds was very tight. The tightness of the range where pathogen virulence increased with the number of spores on the insect body suggests that a very low number of spores enables the death of flies. In this line, Ugine et al. (2005) found, by direct observation of spores on insects, that early-second-instar western flower thrips are susceptible to a strain of *B. bassiana* at a very low dose, approximately 42 conidia per individual. The mechanism of fungal pathogenesis in insects follows several steps: first, spore adhesion, germination, and finally hyphae penetration in the hemocoel. Opposing, the immune response toward *M. anisopliae* in insects reacts in 3 steps: fungus detection, cuticle reaction (e.g., melanisation [Leger et al. 1988a]), cellule immune response, and humoral immune response produced by the fat body (Lu & Leger 2016, Qu & Wang 2018). Hence, the main parameters affecting the dose effect on virulence would be, firstly, where are the spores on the insect body, which translate into how many spores are effective, i.e., germinate and penetrate the insect cuticle. It appears that hairs can prevent adherence to the cuticle and that some body areas are more favourable to adherence (Scholte et al. 2003). In addition, the positioning on the most vulnerable areas of the insect cuticle, such as the intersegmental membrane, affects penetration speed (Leger et al. 1988b, Amnuaykanjanasin et al. 2012) and the thickness of the procuticle is known to be correlated with disease resistance (Charnley 1989). Secondly, insect immune response efficiency can probably be overwhelmed by a minimal number of spores (Rhodes et al. 2018). Microscopic observations of spores on the body of insects would help to better define the ‘minimum’ and ‘optimum’ spore loads. Regarding low inoculation doses, in addition to mortality, sublethal effects might be investigated as they could significantly affect population dynamics by reducing mating competitiveness in males, fecundity and fertility or time for first oviposition in females (Quesada-Moraga et al. 2006b, Dimbi et al. 2013, Duneau et al. 2024).

The three field temperatures tested allowed similar percent of spore germination. The Met69 strain appeared to be tolerant to a wide range of temperatures as a high percentage of spore germination was observed at 20°C (February temperature). In another study on six strains of *M. anisopliae*, Dimbi et al. (2004) found no strain exceeding 70.0% percent germination after 24h at 20°C. Most of their strains grew faster with an increase in temperature from 15 to 25°C, but then slower with an increase in temperature from 25 to 35°C. In our study, the best growth of the Met69 strain within the first 50 h was observed at the hottest temperature 27.7°C (October temperature). In insect-fungus interactions, the latent period of infection and host recovery rate can vary dramatically across and between seasons due to the thermal biology of the host and changes in environmental temperature (Blanford & Thomas 1999). While in some insect species, higher temperatures provide a higher immune response (Adamo & Lovett 2011), here the immune defense of the fly and the mycelium growth appeared to tip in favor of the fungus when temperature increased. Dimbi et al. (2004) found the same until 30°C when inoculating *Ceratitis* species with different strains of *M. anisopliae*. This higher virulence with temperature could be either the result of higher growth of the mycelium or a decrease in the immune response of the fly. As control individuals (not contaminated) died faster when temperature increased, the second option is likely to be the most at play. In addition, the optimal temperature of *M. anisopliae* was found to be between 25 and 32°C with some strains being heat tolerant up to 35°C (Ouedraogo et al. 1997), while the optimal temperature of *B. dorsalis* is between 20 to 30°C (Fiaboe et al. 2021, Rwomushana et al. 2008). Yet, it is also possible that heat shock might be less detrimental to insect immune response than constant high temperatures (Wojda et al. 2009).

The effect of pathogen dose on fly mortality was not influenced by fly sex, but the effect of air temperature on pathogen virulence slightly was. However, this later result should be interpreted with caution due to the unbalanced representation of both sexes in the fertile and sterile populations. Insect sex is suspected to affect their immunity, but it is generally found that, fitting Bateman’s principle (Hangartner et al. 2013), males gain fitness by increasing mating rates, whilst females gain fitness through increased parasite resistance and longevity (Bateman 1948). Luckily, our results did not support this principle in *B. dorsalis*. Indeed, this principle runs counter to what would be beneficial for entomovector technology, since males are generally the ones utilized as vectors and females are inducing the damages.

Formulation, after fungal strain selection, is one of the tools available to modulate the virulence of the pathogen. Adjuvant, either powder or spore coating, can increase the viability of spores by protecting them from ultraviolet radiations (Fernandes et al. 2015), their virulence (Muniz et al. 2020), their adherence on insects (carrying capacity or insect load) (Lu et al. 2020), or their transfer to conspecific using electrostatic properties of the adjuvant (Baxter et al. 2008). Here the mixture with corn starch might only have a ‘dilution’ effect as it decreased pathogen virulence. Thus, it could effectively be used to homogenously inoculate flies at low doses. Low dose led to death but more slowly, which is interesting to increase the transfer duration to conspecifics in the framework of boosted SIT programs. However, this hypothesis requires further testing.

This study provides elements to standardize the evaluation of virulence of entomopathogenic fungal strains against adult fruit flies to optimize auto-dissemination or entomovectoring-based control strategies. Knowledge gathered on low dose efficacy of Met69 against the oriental fruit fly, dilution effect by corn starch adjuvant, and temperature-mediated virulence is of utmost importance to improve control techniques based on auto-inoculation of wild flies or vection by releasing contaminated mass-reared sterile males. The next step should be to conduct field trials that incorporate these findings. In Senegal, the latter technique could be deployed during the off-dry season (from February to May) when fly population is low and concentrated in preferred habitats (e.g., areas with a dense overhead canopy and high relative humidity). This could prevent population outbreaks at the time of mango fruiting, provided that low temperatures (Meats A & Fay 1976) or reproductive arrestment (Clarke et al. 2022) do not reduce fly interactions and, in the end, spore transmission. The approach of this study to calibrate area-wide control strategies, taking into account the host-pathogen interactions and their mediation by abiotic factors from the environment in which they evolve, is a first. It addressed the complexity of biological methods through an integrative approach that can be replicated to develop analogous strategies against other pests.

## Acknowledgments

We thank Real IPM for collaboration and providing the fungal spores.

## Funding

This work was supported by the CEDEAO and AFD (SPRMF), Long-term EU-Africa research and innovation Partnership on food and nutrition security and sustainable Agriculture (LEAP-Agri), Pest-Free Fruit project, in the framework of the European Union’s Horizon 2020 research and innovation program under grant agreement No 727715, and public funds received in the framework of GEOSUD, a project (ANR-10-EQPX-20) of the programme ‘Investissements d’Avenir’ of the French National Research Agency.

## Conflict of interest

The authors declare no conflict of interest.

## Notes

https://doi.org/10.18167/DVN1/X7UTSI

### Competing Interest Statement

The authors have declared no competing interest.

### Summary of Updates

The article has been recommended by PCI Zoology: https://doi.org/10.24072/pci.zool.100282. It has been formatted and the PCI Zoology badge has been added on the 1st page.

