## Appendix for "Dose, temperature and formulation shape *Metarhizium anisopliae* virulence against the oriental fruit fly: lessons for improving on-target control strategies"

### Appendix 1: Effect of the inoculation tube size on the spore load of the fly

Flies were contaminated using cylindrical plastic tubes, lined with velvet. Three dimensions were tested: (6.4 cm*3.5 cm), (7 cm*5 cm) and (8 cm*6 cm). The "standard" tube of 8 cm length and 6 cm diameter (similar to Dimbi et al. 2013) was contaminated at the dose of 40 084 926 spores/cm² of velvet by introducing 0.32 g of dry conidial powder into the tube. The same dose per square centimeter of velvet was respected for each tube size i.e., 0.15 g for a tube of 6.4 cm length and 3.5 cm diameter, and 0.23g for a tube of 7cm length and 5 cm diameter. Five flies were placed in the inoculation tube for 3 min. The flies were then individually immersed in tubes containing 10 ml of distilled water and vortexed for 3 min to unhook the spores on the fly body. A sample of this solution was used for Malassez cell counting. The same process was repeated twice for each tube size which provided 10 replicates i.e., flies, for each tube size. A GLM with a quasiPoisson distribution was used to test the effect of the tube size.

No effect of the tube size was found ($F_{2,27}$= 0.268, P=0.766).

### Appendix 2: Effect of vortexing duration on the number of spores counted on contaminated flies

The "standard" tube of 8 cm in length and 6 cm in diameter (similar to Dimbi et al. 2013) was contaminated at the dose of 40 084 926 spores/cm² of velvet by introducing 0.32 g of dry conidial powder into the tube. Then 5 flies were introduced into the tube and exposed to the conidia for 3 min. Flies were then placed individually in 10 ml tubes of distilled water and vortexed according to three durations: 1 min, 3 min, and 5 min. A sample of this solution was used for Malassez cell counting to determine the number of spores on the fly itself. Two rounds of inoculation were led so that 10 replicates were carried out for each vortexing duration. A GLM with a quasiPoisson distribution was used to test the effect of the vortexing duration, and Tuckey tests were led for vortexing duration pairwise comparisons (package multcomp [Hothorn et al. 2008]).

Vortexing duration significantly affected the number of spores recorded ($F_{2,27}$= 10.618, $P < 0.001$). The difference was significant between 1 and 3 min, while no difference was found between 3 and 5 min. Hence, 3 min was considered sufficient for a reliable spore count.

Table 1: Mean number of spores recorded on the fly according to the vortexing duration.

| **Vortexing duration** | **Mean number (± SE) of spores per fly** |
|---|---|
| **1 min** | 1 608 000 ± 119 970 a |
| **3 min** | 2 580 000 ± 202 232 b |
| **5 min** | 2 336 000 ± 151 813 b |

### Appendix 3. Correspondence between the spore dose in the tube expressed as a density and the fly load

Table 1. Values for the spore doses in the tubes expressed as densities and the corresponding mean fly loads.

| Dose (spore.cm2) | Mean (±SE) fly load |
|---|---|
| 267233 | 4600 ± 783 |
| 400849 | 5200 ± 967 |
| 8016985 | 5600 ± 1101 |
| 40084926 | 193000 ± 26274 |
| 160339703 | 11578000 ± 914936 |
| 320679406 | 23780000 ± 2201396 |
| 641358816 | 23930000 ± 1627319 |

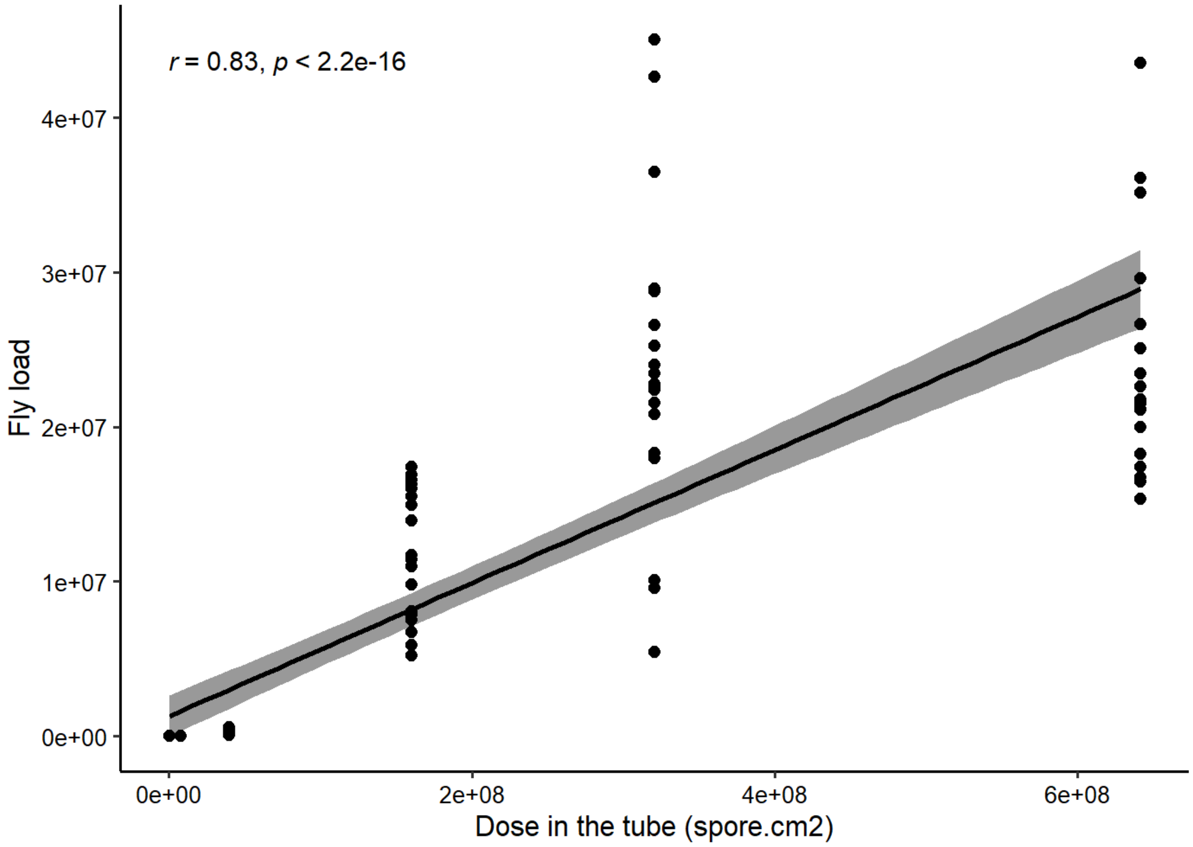


Figure 1. Correlation between the spore doses in the tubes expressed as densities and the fly loads.
